# Natural Gaze Data-Driven Wheelchair

**DOI:** 10.1101/252684

**Authors:** Lou-Ann Raymond, Margherita Piccini, Mahendran Subramanian, Orlov Pavel, Giuseppe Zito, Aldo Faisal

**Author notes:** Equal contribution. L. A. R., M.P., M. S., G.Z., and A. F. are with the Department of Bioengineering, Imperial College London, South Kensington, SW7 2AZ, UK. This work was supported in part by the EPSRC, UK.

## Abstract

Natural eye movements during navigation have long been considered to reflect planning processes and link to user’s future action intention. We investigate here whether natural eye movements during joystick-based navigation of wheel-chairs follow identifiable patterns that are predictive of joystick actions. To place eye movements in context with driving intentions, we combine our eye tracking with a 3D depth camera system, which allows us to identify which eye movements have the floor as gaze target and distinguish them from other non-navigation related eye movements. We find consistent patterns of eye movements on the floor predictive of steering commands issued by the driver in all subjects. Based on this empirical data we developed two gaze decoders using supervised machine learning techniques and enabled each of these drivers to then steer the wheelchair by imagining they were using a joystick to trigger appropriate natural eye movements via motor imagery. We show that all subjects are able to navigate their wheelchair “by eye” learning it within a short time span of minutes. Our work shows that simple gaze-based decoding without need for artificial user interfaces suffices to restore mobility and increasing participation in daily life.

## I. Introduction

THE World Health Organization reports that 2.4 % of the world’s population live with significant difficulties in functioning and require a wheelchair to carry out daily activities. The powered wheelchair is a vital device for many, providing mobility and increasing participation in daily life. However, most of the currently available mechanisms cannot adapt to people with severe disabilities such as high-level spinal cord injury and quadriplegia. The number of spinal cord injuries increases each year, resulting in the urgent need for appropriate technologies to improve quality of life for those suffering from motor disabilities. Due to an increase in wheelchair needs, there has been significant growth in wheelchair research and development.

### A. Human Machine Interface for Wheelchair Control

At present, the need to develop interfaces that provide independent mobility is growing rapidly. Almost all of the readily available control interfaces present several drawbacks and benefits which can be classified depending on the patient autonomy. One of the most common wheelchair control interfaces is the joystick, which is characterised by high information throughput. Furthermore, the joystick provides full patient autonomy. Most are analog systems, and allow the patient to adjust wheelchair speed. But, this requires considerable skills however, and is hence not suitable for severely disabled patients. Another low-cost device that has seen some adoption is the sip/puff interface, which provides directional commands by inhaling (sipping) and exhaling (puffing) through a straw, which in turn activates a pneumatic switch or pressure sensor. Unfortunately however, the response of the interface is slow and the tubing requires frequent cleaning [Jeonghee Kim 2013].

Brain Machine Interfaces provide hands-free wheelchair navigation and ease-of-use for those with limited upper limb mobility. Brain Computer Interfaces use invasive and noninvasive electrodes that acquire biological signals. One of these interfaces, EEG (electroencephalography), uses electrodes placed on the patient’s scalp to measure the electrical activity of the brain in order to control the wheelchair. The user is trained to voluntarily modulate the EEG by executing mental imagery corresponding to each of the steering commands [Ferran G et al., 2008]. Limitations include very low information throughput and transfer rate, making the system insufficient to quickly react to dynamic changes during wheelchair navigation [Brice Rebsamen et al., 2010]. In [Abbott and Faisal 2012], it is shown that the decoding rate of brain information (information throughput) ranged from 0.5 to 7 bits per second, and that the wheelchair controller requires a higher rate of at least 15.3 bits per second. Furthermore, it presents a high sensitivity to noise, related to the acquisition of brain signals [Richard C Simpson 2005].

### B. Gaze Control for Wheelchair Driving

The human visual system is a multifaceted optical scheme in which the image is converted into a sequence of electrical signals, and is subsequently communicated to the brain through complex neural schema [Land, M. F]. Eye gazing plays a significant role from infancy onwards, and plays a vital part in nonverbal communication [Farroni, T; Fullwood C; Mayer K; Estrada CA]. Moreover, eye movements are highly interrelated with motor intents, and are often retained by humans with severe motor deficiencies [Ruofei Xu]. Applied physics and modem electronics make real-time eye gaze tracking a possibility. On-chip charged coupled devices with digital signal microprocessors have significantly increased the usability, accuracy, and speed of eye-tracking technology while decreasing the cost [Duchowski, A]. Human-computer interaction is the most desired use of eye tracking, particularly for gaze-contingent communication [Duchowski, A. T. (2002); Orlov, P. A., & Apraksin, N. (2015)], and has various applications in biomedical engineering [Ktena, S. I et al.; Eivazi, S et al.]. Gaze driven systems provide hands-free control of robotic devices [Tostado, P.M et al., 2016; Dziemian, S., 2016]. Eye gaze tracking also allows us to control machines such as human-machine interfaces for semi-autonomous vehicles. One primary application is the use of this system of communication for steering a wheelchair [Martin Tall et al., 2009; Erik Wastlund et al., 2010; Gunda Gautam et al., 2014]. For instance, semi-autonomous and autonomous control of a wheelchair was developed by Ruofei Xu et al. to assist users with severe motor neuron disorders [Ruofei Xu]. In its standalone mode, the user can select a position on a pre-built map and the vehicle will navigate to the desired location. Even though obstacle localization and avoidance algorithms are not refined enough to achieve acceptable accuracy with this setup [Ruofei Xu], the eye-tracker system, in contrary, provides the ability to move independently and safely within the surrounding environment with useful hands-free input. This helpful function can contribute to the user’s satisfaction and learning motivation [Lisbeth M Nilsson and Per J Nyberg 2003].

The eye-tracking systems provide information about the eye movements and gaze location that is not enough for the direct control [Andrew Duchowski 2007]. Imperative difficulty being every gaze should not turn to goal - ‘Midas touch’ problem. Conventional techniques consist of on-screen buttons on a video scene feed, with the user gazing at the buttons to navigate [Wästlund, E 2010]. Other possible iteration techniques include dwell time for activating directions to avoid unintentional movement of the wheelchair [Arai, K]. This method of superimposition was well understood by the subjects and was able to control the powered wheelchair using eye movements within the environment. The screen-based interface requires shifting of attention between the physical navigation environment and the computer display, which can be difficult and might cause cybersickness. Other user interface approaches for gaze control of powered wheelchairs include primary mobile systems such as the optic mouse, on-screen buttons, commercial systems (SMI iViewX RED and Tobii 1750), and webcam based systems [Martin Tall et al., 2009]. A wheelchair steering controlled system based on eye movements allowing the user to move in any desirable direction has been demonstrated too [Gunda Gautam et al., 2014]. However, our earlier research and other studies tried to provide a gaze-driven control system based on natural gaze behavior [Sofia Ira Ketna 2015]. First, natural eye-movements during keyboard control were recorded, followed by gaze data while using each of the interfaces for the virtual wheelchair keyboard control. Such demonstrations show the necessity of using natural gaze behavior as signal commands to control wheelchair navigation [Christian Bartolein et al., 2008]. Most of the aforementioned studies propose eye-based wheelchair control interfaces, which often require more effort by the user and lead to abnormal gaze behavior. It was observed that the visual information was used in a feed-forward manner during the approach phase, but not while stepping over the obstacle.

Task and context determine gaze allocation, and it is not guided according to a random or a saliency model [Yarbus, A. 1967, Constantin A. Rothkopf, 2007]. Marius’t et al. observed that gaze allocation in free viewing condition differs from that under laboratory condition. Thus, factors guiding human eye movements under natural conditions need investigation [Marius’t Hart B, 2009]. One possible factor is compensatory movement, i.e., eye movements go to the left when the head moves to the right. During walking tasks, terrain regularity results in differences in head orientation and gaze behavior, specifically in the vertical direction, and increases in attention while walking on complex surfaces [Marius’t Hart B, 2012]. During natural walking or wheelchair driving the body movements also compensate for gaze instability. Previous studies show that eye-movements play a part compensatory role, and about 20% of eye-movements act synergistically with head movements to direct gaze [Einhäuser W, 2007; Wolfgang Einhäuser, 2009].

A particular environmental feature can be utilized as a visual pivot, for example, the opening of a door, and the safest navigation, defined as the locomotion, is characterized by the fixations toward the door opening. This voluntary fixation is likely related to object detection purpose [Bernard Marius ‘t Hart, 2013]. Takahiro et al., analyzed the location of obsession and the duration of locomotion before the aperture crossing with four forms of movement, including regular walking, walking with a bar, blocked shoulder rotation and finally locomotion with a wheelchair [Takahiro Higuchi, Michael E Cinelli 2013]. Due to the influence of unfamiliarity with the system, another study focused on the enhancement in space estimation ability for movement with familiar and unfamiliar wheelchair users [Takahiro Huguchi et al., 2009]. Advanced navigation techniques mediated by natural free view achieved through sophisticated algorithms would be the ideal next step in this field [Ktena, S.I].

Herein, we investigate people’s gaze pattern while driving a wheelchair using an RGB-D (Depth) camera. This has proven essential for developing an eye-based control interface which retains the driver’s natural gaze behavior while controlling the wheelchair with the eyes. In particular, we are interested in investigating whether the user’s visual attention is focused on the floor while following a specific path and whether this behavior is a result of the complexity of the environment. Also, we search for correspondence between people’s gaze patterns and motor intentions. To achieve this goal, we firstly built a wheelchair with an eye-tracker and RGB-D camera, and aligned RGB-D data with gaze data. Secondly, we collected gaze points according to driving actions and floor position. Finally, we built two natural gaze data-contingent driver interfaces.

## II. Method

This section explains the methods deployed for the alignment, segmentation, estimation, visualization, and accuracy estimation of the gaze position using a depth camera and eye-tracker.

### A. Eye tracker calibration on the wheelchair

A 60fps eye-tracker was used for this study. We performed basic 2D eye-tracker calibration included in a commercial eyetracking software package. The remote eye-tracker provides gaze data mapped to the 2D video frame - our strategy by contrast involves analysis of gaze data in a physical 3D space. To obtain the depth information of the surrounding environment we used an RGB-D camera. By combining the RGB-D camera and eye-tracker information throughput, we generated a 3D gaze dataset. Before expressing this gaze in 3D, the 2D eye-tracker video frame was aligned with a RGB-D frame by applying an affine transformation between the two coordinate systems. The alignment method consists of several steps. First, 10 OptiTrack reflective markers were placed on the floor in front of the wheelchair (Fig. 1) in such a way as to be detectable by both the RGB-D camera and the eye-tracker.

**Fig. 1.**
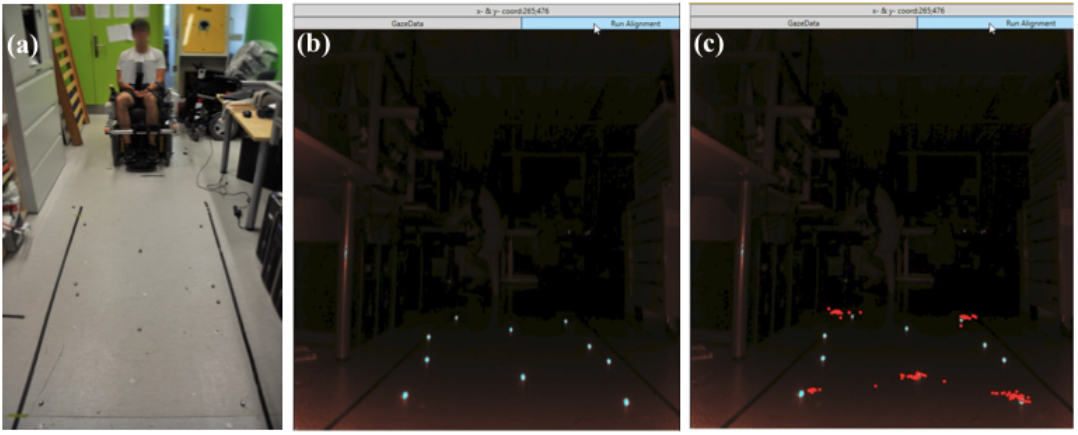
(a): Alignment setup. OptiTrack markers in the working space can be seen in front of the wheelchair, (b): Alignment of the 2D eye-tracker video frame with a RGB-D frame through affine transformation between the two coordinate systems. (c): Qualitative analysis of accuracy of the affine transformation applied to the eye tracker and the RGB-D camera coordinate systems.

The RGB-D camera image was mirrored to facilitate the visualization of the surrounding environment. The top right corner of the image is, therefore, the origin of the infrared frame reference with x=0 and y=0. The user is asked to look at each of these markers consistently. The origin of the eye tracker frame reference corresponds to the top left corner of the panel with x=0 and y=0. In both systems, the pixels express coordinates. The alignment between the two coordinate systems was applied without any need to add a scaling transformation to enlarge or shrink the objects in all directions.

Once the gaze data and the image coordinates are normalized, an affine transformation is applied to express the gaze data in the infrared sensor frame of reference. The affine transform matrix ‘A’ allow the application to each gaze point [Helmuth Spath 2004]:

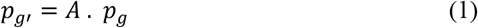

where Pg’ corresponds to the definition of each gaze point in the new reference system (infrared sensor) and Pg corresponds to the gaze point in the eye tracker system. The matrix ‘A’ is defined in the least squares fashion. The least square estimation of A is:

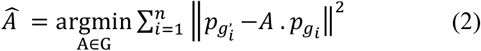

where G is an affine group, then (1) can be expressed as:

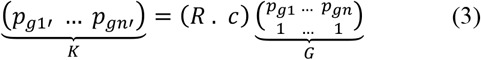

where, for our application, K is the ‘RGB-D camera’ data, and G is a matrix composed of the ‘Gaze’ data. ‘n’ is the number of points, ‘R’ is the rotation matrix of the affine transformation and ‘c’ corresponds to the translation. Both ‘R’ and ‘c’ included in ‘A’. The ten markers coordinate expressed in the eye-tracker reference frame, and the RGB-D camera reference frame are respectively the values of the array G and the matrix K. Finally, the least squares estimation of A is given by:

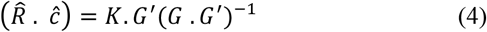

where G’ is the transpose of G. Once and are defined, these terms used in (1) in matrix ‘A’ and each gaze point, expressed in the infrared sensor reference frame. Fig. 1 represents the results obtained by applying the affine transformation, allows verifying the accuracy of the alignment qualitatively. The red points correspond to each new gaze data point expressed in realtime in the reference frame of the infrared sensor.

### B. Floor Detection using the RGB-D camera

To separate gaze data on floor-oriented and no-floor oriented gaze points, we detected the floor’s position with a built-in function of the RGB-D camera. This function estimates the coefficients of the floor plane in standard hessian form with the normalized plane equation:

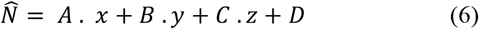

where A, B, and C are the components of a unit vector corresponding to the normal of the plane and D is the distance from the origin of the camera to the plane (fig. 2).

**Fig. 2.**
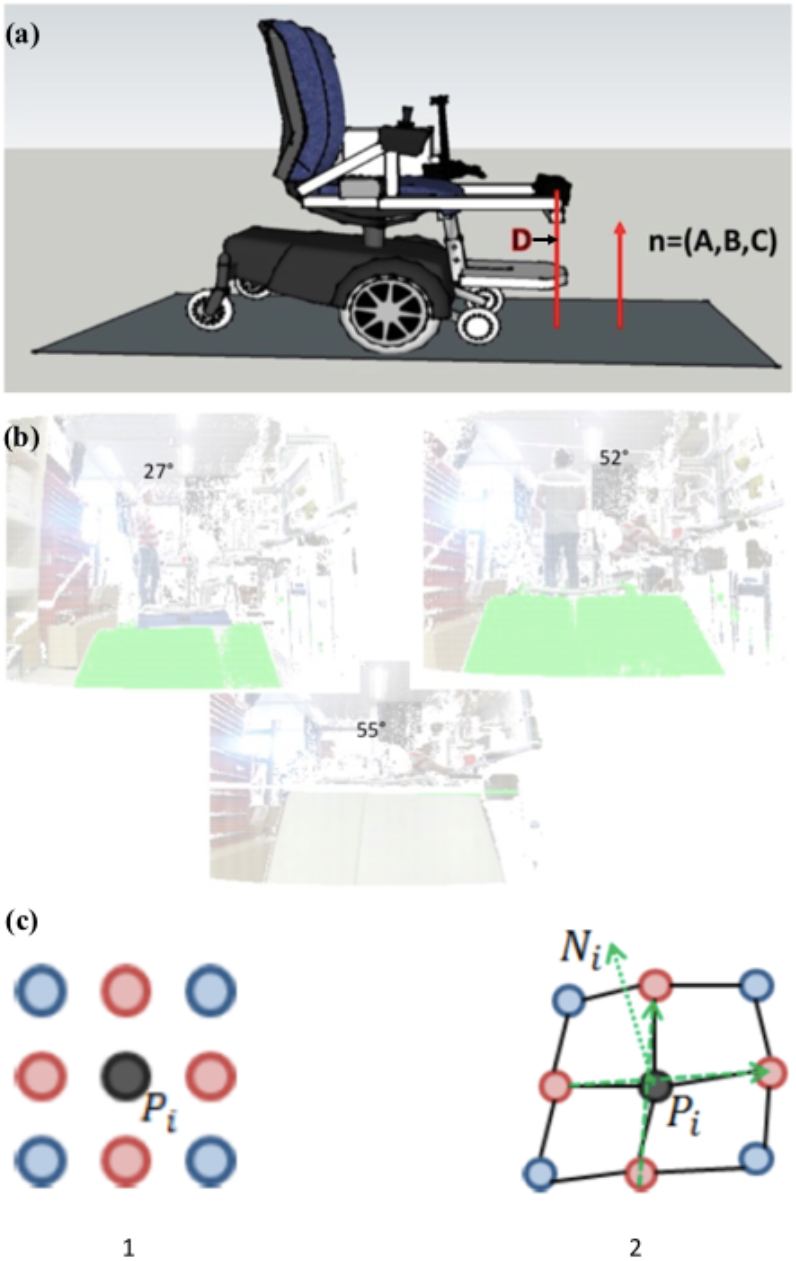
(a): Illustration of the floor plane equation estimation, D is the distance from the RGB-D camera to the floor and n is the normal to the floor, (b): Tilted floor detection, visualization of the detection of tilted floor with the first algorithm using the built-in function. (c) Local normal vectors computation: 1) Selection of the neighbours of point Pi on the depth map. 2) Vectors up-down and left-right in the 3D point cloud [H. W. Yoo et al., 2013].

This floor detection algorithm has a low computational cost but is not robust. Using this algorithm, only floors tilted under 55° can be detected. A clipping plane renders only specific portions of the scene and excludes some parts of a scene’s geometry. It is sensitive to the RGB-D camera vibrations leading to some computational errors such as incorrect parameters values. In fact, the structure of the built-in function requires that the camera is parallel to the ground level. If the camera is tilted, the parameter D, the distance plane-camera, is null and the other parameters are wrongly estimated. The vibrations of the system while the wheelchair is in motion makes the camera tilted and, due to the lack of the algorithm robustness, fail to evaluate the floor as a clipping plane. A solution to solve this vibration issue is to attach the RGB-D camera firmly to the wheelchair via a steel frame, fixed to a stable structure above the chair’s wheels. This fixation strategy improves the robustness of the floor detection algorithm substantially. Furthermore, this algorithm can detect a tilted floor as shown in Fig. 2. This floor detection strategy was used in the data analysis to verify the gaze pattern and compares between floor/non-floor positions. It is also implemented in the wheelchair controller interface to enable the distinction between motion and no motion following the results obtained after the data analysis presented in the results.

### C. Wheelchair setup evaluation

We conducted a use-case study to test the accuracy of the 3D gaze data estimation by the combination of eye tracker and depth camera. For the analysis of the significance between groups of data, we performed a one-way Analysis of Variance (ANOVA) followed by Tukey Honest Significant Difference (HSD) for posthoc comparisons test; p-value > 0.05 considered as not significant. Comparison data for the tests are represented in bar graphs with the standard deviation of no less than N = 3 to 11 subjects.

To test the 3D gaze data accuracy 7 healthy subjects (22 - 30 years old) with normal or corrected to normal vision were asked to fix at the markers on the floor in different patterns. The accuracy based on the comparison between the 3D coordinates of the gaze point on a particular location on the floor and the 3D coordinates of this location obtained using the OptiTrack Motion Capture cameras. The OptiTrack cameras have a precision in the order of millimeters and provide accurate 3D world space coordinates of the location of the reflective OptiTrack marker. Three cameras have been used and calibrated. They were facing the experimental setup including the wheelchair as shown in Fig. 3.

**Fig. 3.**
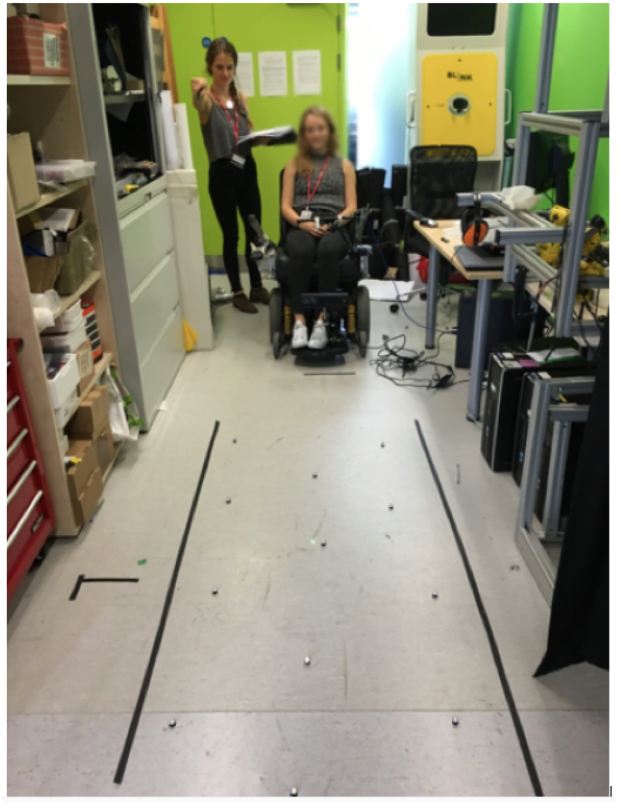
Test of the 3D Gaze Data Accuracy. The technician is using a laser to indicate 10 different patterns to the user.

The RGB-D camera space and the OptiTrack system did not have the same origin of the reference frame. The source of the RGB-D sensor was first detected using the reflective markers and expressed following the origin of the OptiTrack system. For the translation that had to be applied to the final 3D gaze data, in order to compare their accuracy with the OptiTrack measurements. Once the RGB-D camera origin is detected, 15 markers placed on the floor facing the wheelchair, and their coordinates using the OptiTrack cameras, were saved to a file. We then obtained the 3D gaze data at these 15 marker locations using the current setup, following three steps:

1. A user profile was generated using the 2D eye-tracker calibration data.
2. The 2D alignment between the RGB-D camera infrared sensor and the 2D gaze data was applied using ten markers.
3. The 15 markers utilized for the OptiTrack data were placed on the ground. Subjects were asked to look at the 15 markers following different patterns indicated by a technician using a laser as shown in the fig. 3.

We recorded 3D gaze data, and the translation between the two systems origin was applied to each of these data points to estimate the accuracy, the Euclidean error between the 3D gaze data, and the OptiTrack data for each marker calculated. The results obtained and show in Figure 4. highlight the mean Euclidean error over the 15 OptiTrack markers locations for the ten different patterns. By averaging across the subject, the estimated 3D gaze data acquired had a Euclidean error of 16.7 ± 1.2cm. Impreciseness between the ground truth and the 3D estimated data was insignificant. This value was considered to be accurate enough for our application.

**Fig. 4.**
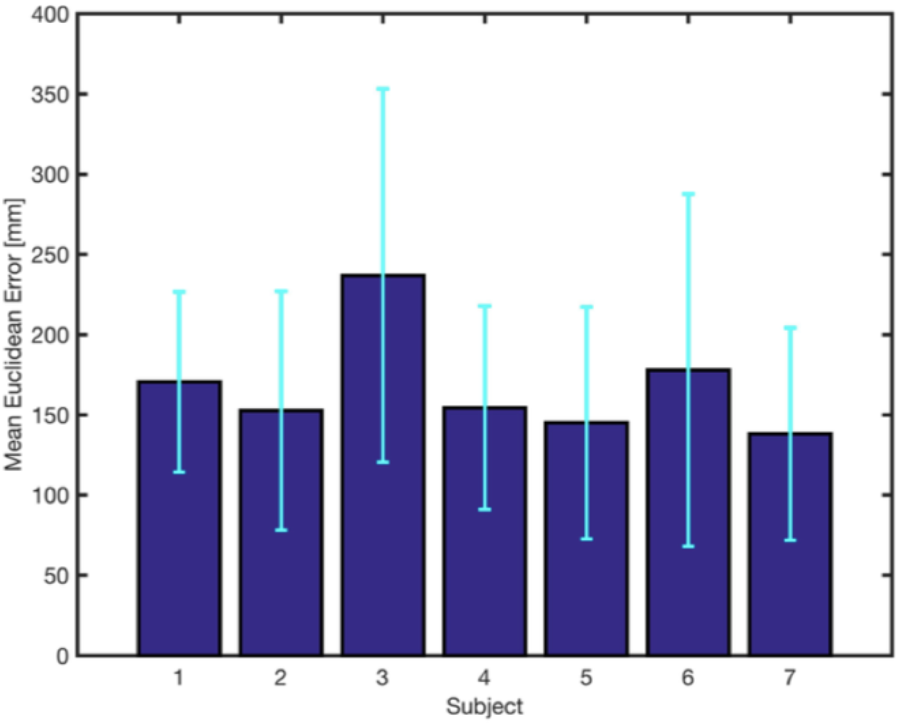
Mean Euclidean error of 3D gaze estimation for each subject. Error bars represent the standard deviation obtained comparing the ground truth with the estimated.

We observed that the 3D gaze data obtained at the extremities of the original working space, being limited by both the RGB-D camera and the eye-tracker, were wrongly estimated and presented significant error. The RGB-D camera/eye-tracker system was inefficient at these locations due to the combination of their field of view limitations. The workspace is then limited to the range of -1 to 1 meters laterally from the RGB-D camera space origin, and 0.5 meters to 2.5 meters in depth from the origin. These ranges limited the number of valid gaze data points usable for the investigation of natural gaze behavior. It therefore limits some valid gaze points accessible for the controller interfaces to navigate safely and rigorously in the surrounding environment with the eye-based wheelchair.

## III. Natural eye movements study during navigation

### A. Experiment design

We conducted a study to collect the 3D gaze data and then used it for the decoder development as training data. 8 healthy subjects (22 - 30 years old) with normal or corrected-to-normal vision were involved in the study. All of them were naive to the use of a wheelchair. Each experiment trial started with the calibration of the eye tracker, followed by the alignment procedure to calibrate the subject, which lasts for roughly 20 minutes. The experiment consisted of three different tasks, where the participant had to navigate with the wheelchair in three different environments freely, using the joystick. In task one, the subject had to go back and forth in a hallway free from obstacles. In task two, the participant was asked to navigate in a room with three chairs and moved around these obstacles for about 5 minutes. In task three, the subject was placed in a more complex environment, cluttered with several tables and chairs, where he/she had to navigate for about 5 minutes again. The 3D gaze data were recorded during all chair movement, as well as during the time between tasks, when the participant moved from one place to the other.

The entire experiment (including calibration, alignment, and recording) lasted 1 hour on average. Therefore, to monitor potential calibration lost during such a lengthy period, a qualitative check of the calibration procedure was conducted before and after each task, where the subject was asked to fixate on five markers on the floor, and the calibration error qualitatively monitored in real time through the video recording. During all the recording, the participant was asked to sit in a comfortable position and to not rotate the upper part of their body. The upper body constraint was necessary due to the system Field Of View (FOV) limitations. However, small head movements were allowed inside the eye tracking box. A belt was used to keep the subject in the same position while he was calibrated to impose the body limitation.

### B. Preprocessing and data analysis

Before proceeding with the analysis, we conducted preprocessing on the raw data. The joystick values did not synchronize with the 3D gaze point data and plane equation coefficients because the values came from two different systems. Moreover, the gaze data were acquired with a nonuniform frame rate, because no values were saved when the eyes of the driver were not detected, either because the user was blinking, or because the user moved outside of the eye tracking box. Therefore, to synchronize the data, the signals were interpolated through the shape-preserving piecewise cubic, named PCHIP (Piecewise Cubic Hermite Interpolating Polynomial) and were re-sampled at a frequency of 50Hz.

With the data synchronised, the plane equation was used to extract the floor / non-floor information, to classify the gaze point of the user. Given a b c and d, parameters of the plane equation (*ax* + *by* + *cz* = *d*) provided by the floor detection algorithm, the distance between the gaze point and the floor plane was computed, using the equation:

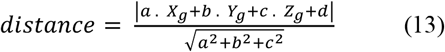

where *X_g_, Y_g_, Z_g_* are the 3D estimates of the gaze point. Given the gaze-floor distance, a threshold of 10 cm was used to classify a point as belonging to the floor, or not. The limit has been selected taking into account the mean Euclidean error of the 3D gaze data estimation, which could not allow a more precise estimate with a lower threshold.

In the first part of our analysis, people’s percentage of gaze points on the floor were investigated, to determine whether or not people look at the floor while driving a wheelchair, and how this behavior changes with the complexity of the environment.

For each subject, the percentage of gaze point on the floor, in the FOV, was computed during the execution of three different tasks. In Fig. 5(A), a common pattern was observed between a group of participants (subject 1, subject 6, subject 7, subject 8). The percentage of gaze fixations on the floor is smallest during task one (free hallway), but increases during task two (room with three chairs), and reaches a maximum during task three (the highly complex environment). A similar pattern can be observed in subjects 3 and 4, who directed the eyes toward the floor more during task two and task three, than during task one. Different gaze behaviours have been identified in subject 5 and subject 2. Overall, a general pattern results from averaging over the subjects (fig. 5(B)). This pattern shows that, overall, the participants directed the fixations toward the floor, more during task two (59,7% ± 7,2% SD) and task three (60,7% ± 14,9% SD) than in task one (45,7% ± 18,2% SD).

**Fig. 5.**
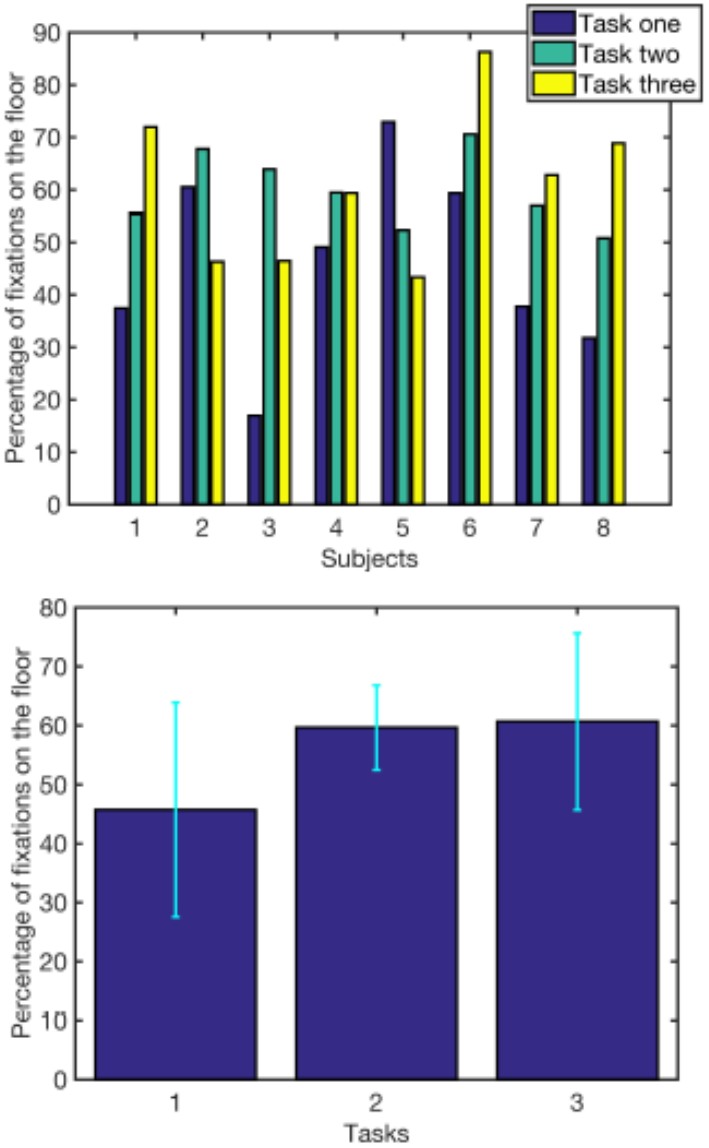
Percentage of gaze points on the floor during Tasks solving (a) Comparisons between different subjects, (b) Mean across 8 subjects.

The result showed a significant main effect (F(3,21) = 3.25, p = 0.04). The mean percentage of gaze points on the floor is lower for task one than for task two (p-value= 0.1) and task three (p-value=0.075). However, the power of the performed statistics is not sufficient to reach significance.

To efficiently design the wheelchair control interface it is essential to distinguish between driving and idle state. Total Recording (no tasks separation), for each participant the conditional probabilities of motion/idle, given the 3D gaze point on the floor and outside of the floor, have been computed with Bayes’ Theorem and are shown in fig. 6(B) For all the subjects, P(Motion | Gaze on the floor) is significantly higher (0:85 ± 0:08 SD) than the P(No Motion | Gaze on the floor).

**Fig. 6.**
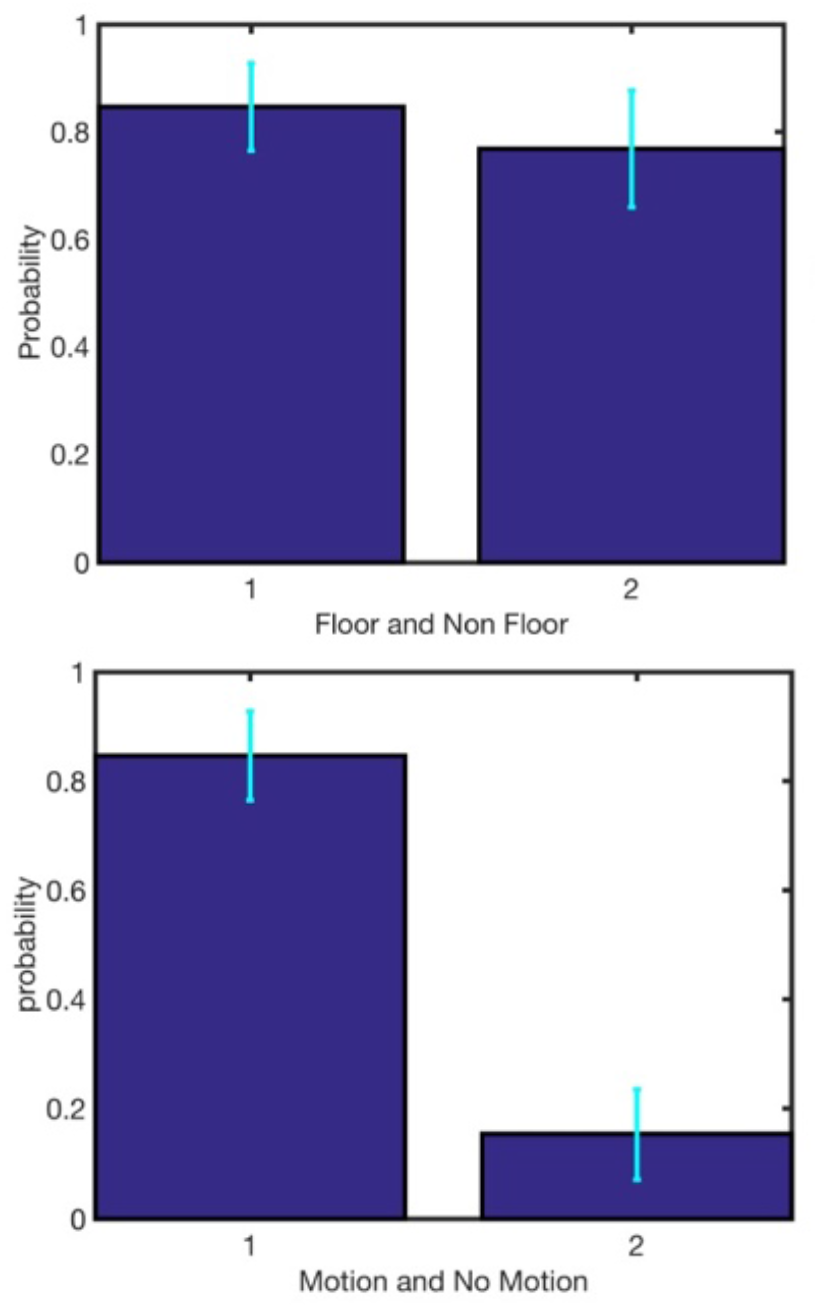
(a) Comparison between the conditional probability of motion given gaze point on the floor and not on the floor. Bar 1 is *P(Motion* | *Floor*); bar 2 is *P(Motion* | *Non Floor*). Mean across 8 subjects (error bars represent standard deviation). (b) Conditional probability of motion or no motion, given gaze point on the floor. Bar 1 is *P(Motion* | *Gaze on floor*); bar 2 is *P(No Motion* | *Gaze on floor).* Mean across 8 subjects (error bars represent standard deviation).

We then measured the P(Motion | Non-Floor) and assumed, for this analysis, that the gaze points outside of the FOV do not fall on the floor. This was necessary given the reduced size of our FOV. Given people’s natural behavior, the probabilities of moving are high, both when the gaze point falls on the floor and when it falls outside of the floor. However, the probability that the driver intends to move is higher in the first case than in the second, as can be seen in Fig. 6(A) The results of related samples t-test show that there is a difference between P(Motion | Floor) and P(Motion | Non-floor)(t(7) = -4.006, p = .005). This confirms the probability that the driver intends to move is higher when the gaze point falls on the floor than when the gaze point falls not on the floor or outside the FOV.

In the second part of the analysis, probability space maps were computed based on Bayes Theorem, to identify regions where the fixations of the driver are likely to fall during a particular motion state (e.g., forward, backward, right, left motion). In each location in the space, these probability maps contain the probability that the driver, given a fixation in that particular location, was in a specific motion state. Different probability maps are computed for gaze distribution on the floor and not on the floor. For a specific state of motion S_i_, the probability space map is computed through a fully-expanded expression of Bayes’ Theorem:

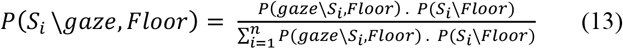

Where

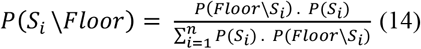

The variable Floor, in (13) and (14) represents a categorical variable, which can be either floor or non-floor.

To apply these equations to the data, the 3D gaze values are divided into two different sample spaces: floor or non-floor data. Each of these two groups is then partitioned in a collection of n subsets, based on the joystick values. Each subset represents, therefore, a particular motion state *S_i_ (i = 1, … n*). The range of x and y joystick coordinates, used to identify different motion states, was manually selected by measuring the joystick values when the wheelchair was stationary and when the joystick was moved to the 4 extreme positions. Using the partitions resulted from dividing the data in the different range of movements, *P(gaze*|*S_i_, Floor*) is computed using the relative occurrence of gazes in each location of the space, divided into cubes of about 5 x 5 x 5 cm3.

Furthermore, *P(Floor|S_i_*) is the probability that the user was/was not looking at the floor while being in the state *S_i_* and *P(S_i_*) is the probability of that particular state. The posterior probability *P(S_i_|gaze, Floor*) is; therefore, a four-dimensional histogram resulted from dividing the x, y and z dimension of FOV in bins of about 5 cm. The value of the histogram in the bins represents the probability of the state, given a gaze point in that particular location. However, the range of y values is minimal, due to the FOV limitation along the vertical axis (fig. 7). Therefore, the probabilities in the floor and non-floor cases are averaged over the vertical dimension and are visualized through heat maps. Mean filtering using a 3 x 3 square kernel 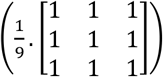 is used to smooth the data. Fig. 8. shows heat maps that allowed us to analyze where people look during different motion states qualitatively.

**Fig. 7.**
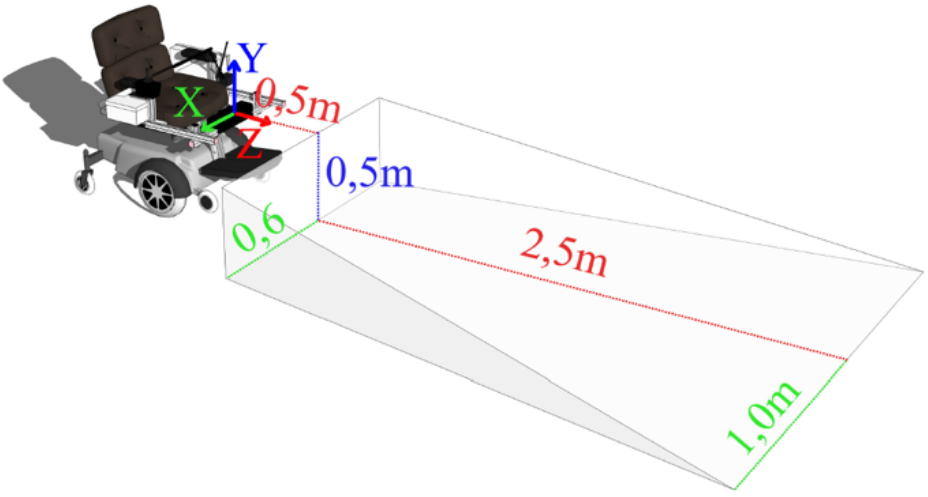
System Field of View

**Fig. 8.**
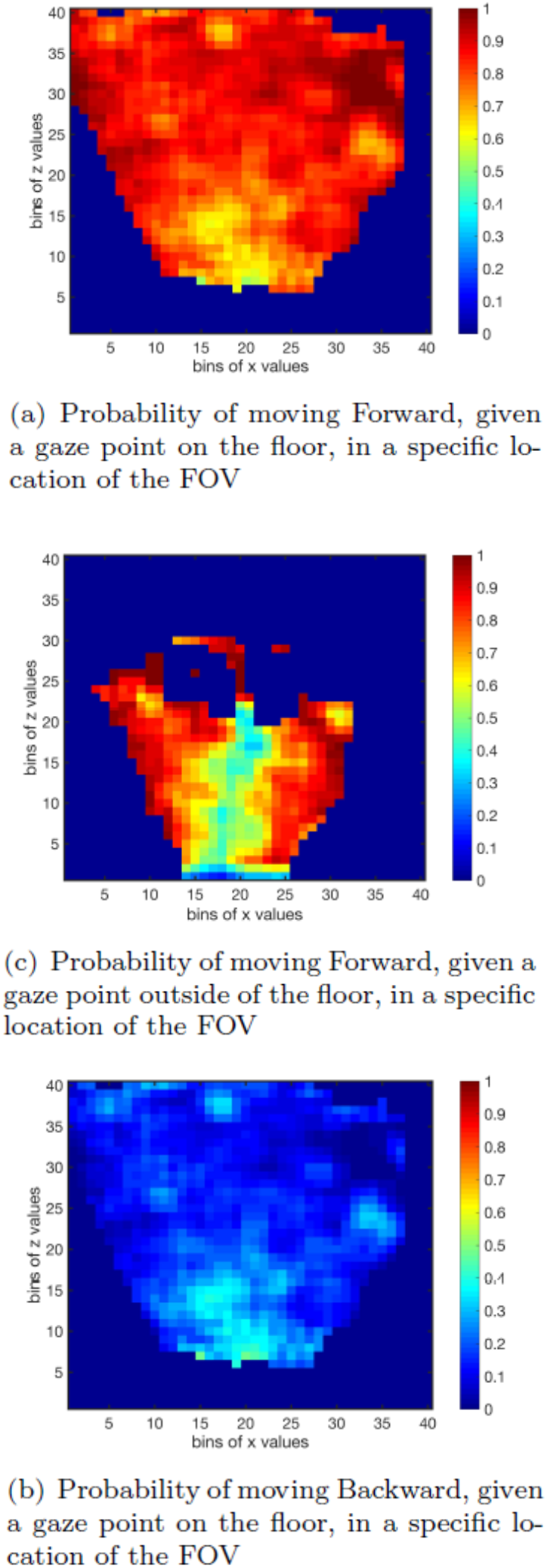
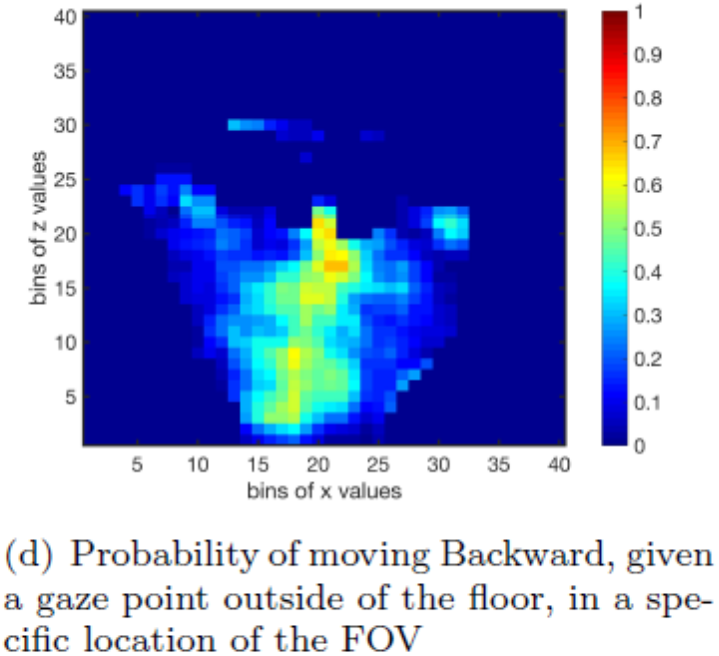
Heat maps for the Forward and Backward motion. Color bars represent the probability of moving in a specific direction, given gaze points on specific region in the field of view.

The conditional probabilities heat maps are shown in Fig. 8. In the heat maps, the number of bins is chosen so that each bin corresponds to about 5 cm in the FOV. Note that the z dimension represents the depth (distance from the wheelchair), increasing with higher bin number. The x dimension represents the width of the FOV. Bin 0 corresponds to the left corner, and bin 40 corresponds to the right corner. First, differences in the gaze pattern during forward and backward motion are analyzed, in the cases of gaze distributions on the floor and outside of the floor. A forward movement is considered when the wheelchair moves forward (including forward right and forward left).

In the heat maps for the forward and backward case (fig. 8), given a gaze point in the FOV, the probability that the user was moving forward (a and c) is higher than the probability that the user was moving backward (b and d). This is because the participants spent naturally more time moving forwards than backwards.

However, when we analyzed the heat maps for gaze distribution on the floor, we observed that when the gaze point falls further from the wheelchair (large z bins), there are higher probabilities of intending to move forward than when the gaze point falls closer to the wheelchair (a). In contrast, for the backward case, we observe that when the gaze points fall close to the wheelchair, there is a higher probability of moving backward than when the gaze point falls further from the wheelchair(b). Such a pattern is not visible when the people’s gaze distributions not on the floor are considered (c and d). Therefore, we can conclude that, when the users look at the floor, they prefer to gaze at the space closer to the wheelchair while driving backward, further from the wheelchair while driving forward. Similarly, the second group of the heat map is computed to analyze where the participant looked while moving to the right and to the left (without taking into account whether the movement was in forward or backward direction). In this case, the participants were likely to look at the floor on the right or left side of the FOV, respectively, while moving rightward or leftward, respectively (fig. 9(a,b)). This pattern was confirmed, when the gaze data considered did not fall on the floor (fig. 9(c,d)).

**Fig. 9.**
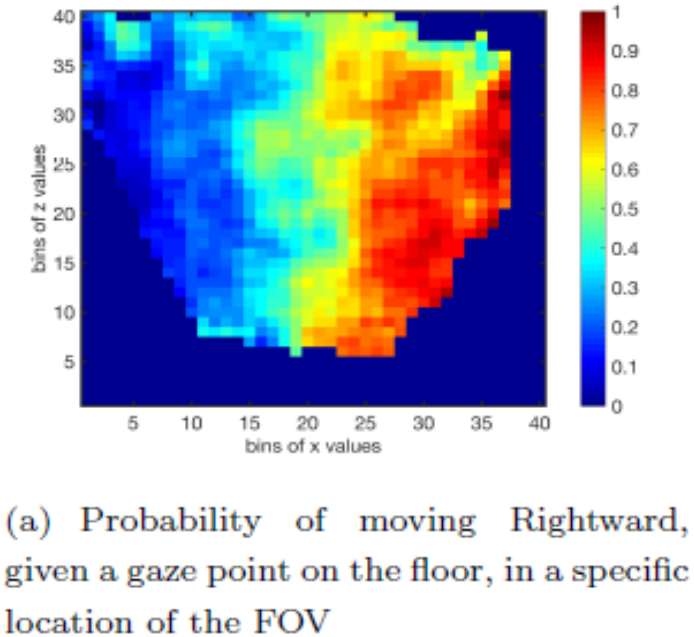
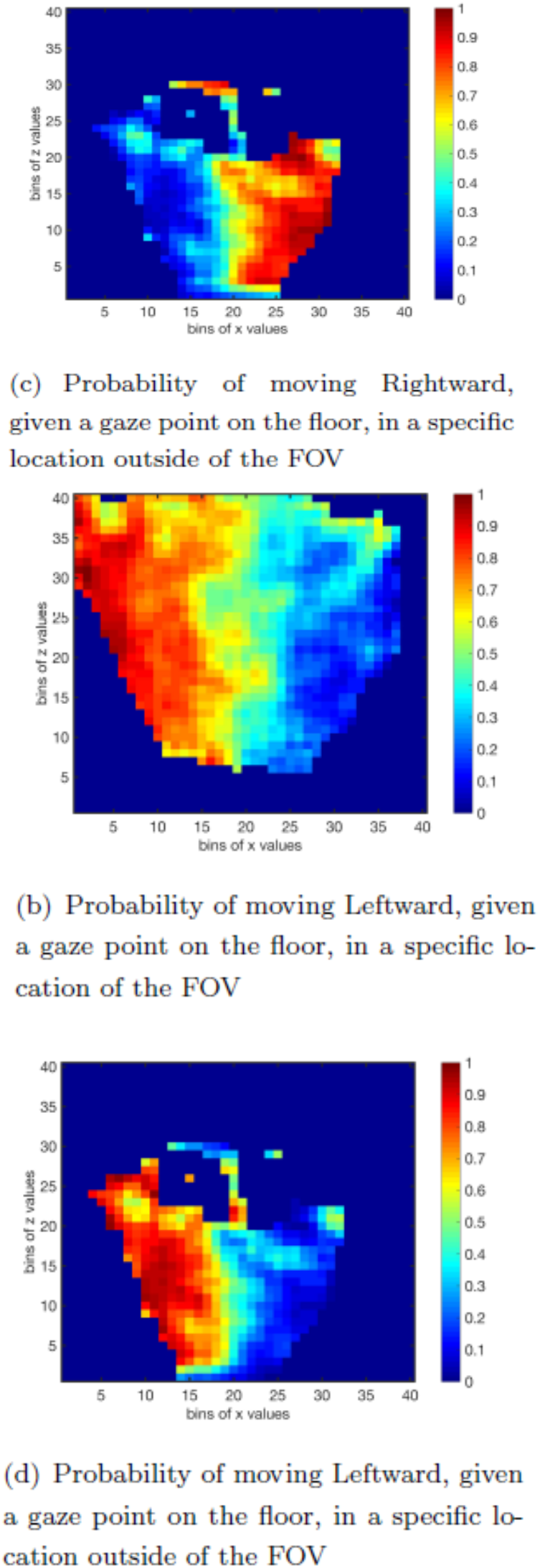
Heat maps for the Rightward and Leftward motion. Color bars represent the probability of moving in a specific direction, given gaze points on specific region in the field of view.

Gaze patterns for four different directions of motion have been compared (see Fig. 10): Forward Left, Forward Right, Backward Left and Backward Right. If we observe the heat maps for the movement in the forward direction (including Forward Right and Forward Left) (fig. 10(a, b, c, d)), we can see that the previously discussed pattern are still visible. For example, there is a preference for the users to look at the right side of the FOV, further from the wheelchair, while moving forward right. However, when we consider the heat maps for the motion in the backward direction (including Backward Left and Backward Right) (fig. 10(e, f, g, h)), a pattern is not visible anymore and no real conclusion can be drawn. These results showed that the apparent trend shown in fig.9 is mainly due to the forward motion. This leads to the conclusion that people tend to look in the direction they are moving, during the forward state, yet no clear difference exists between looking right and left during the backward state.

**Fig. 10.**
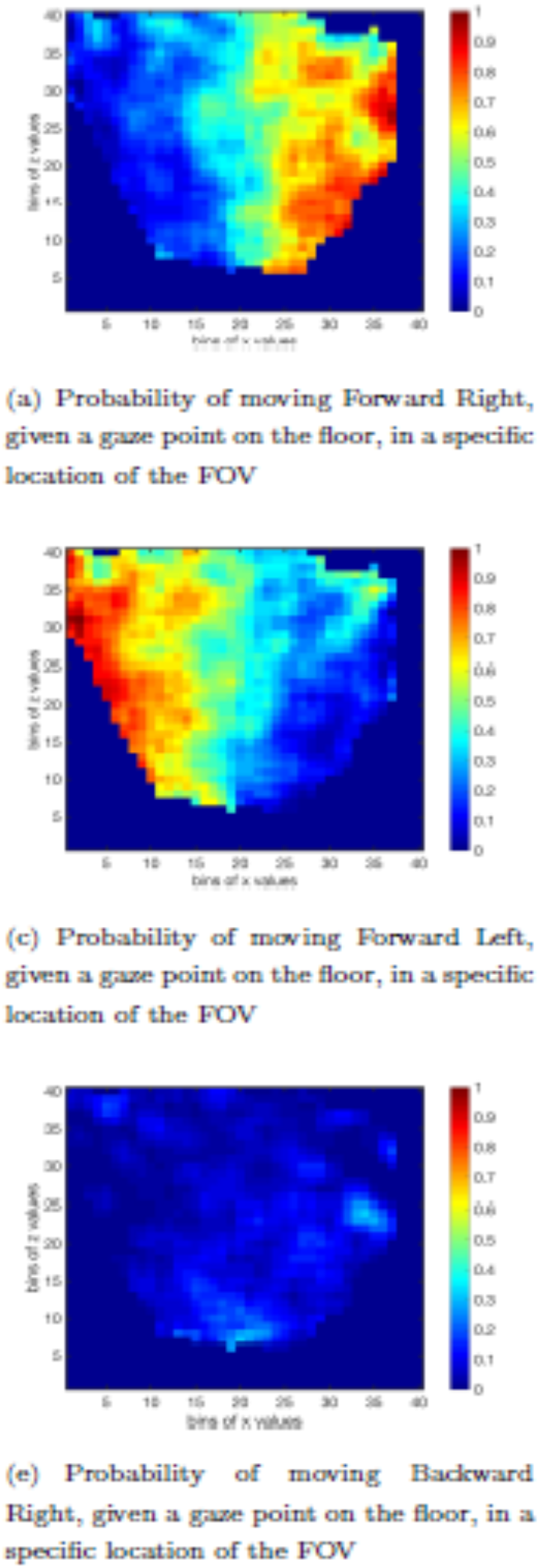
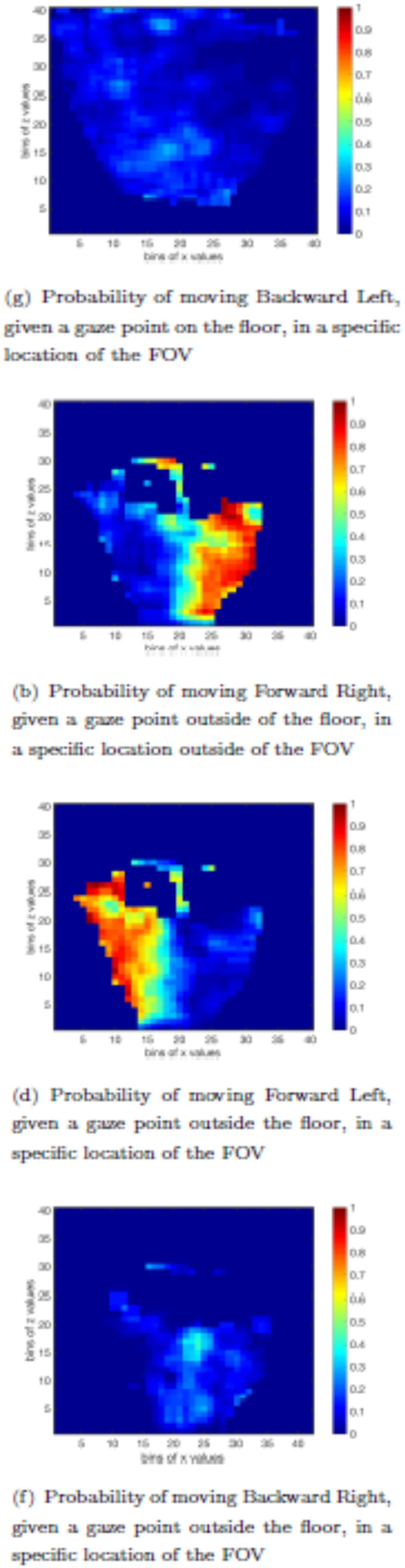
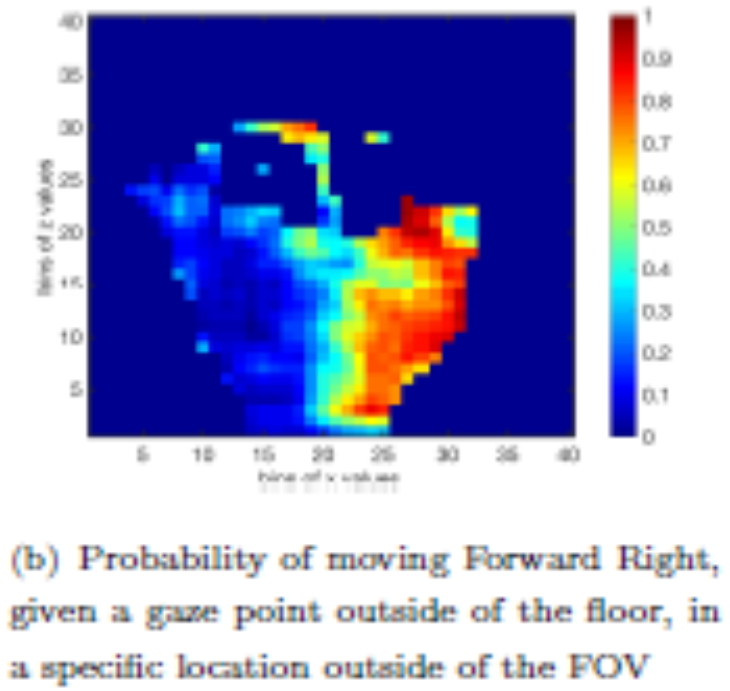
Heat maps for Forward Right – Forward Left – Backward Right – Backward Left motion. Color bars represent the probability of moving in a specific direction, given gaze points on specific region in the field of view.

Based on these results, two different interfaces have been proposed to translate the gaze data into bit values in the range of the National Instruments (NI) controller, to generate the desired wheelchair motion. Both the interfaces use the x and the z coordinates of the 3D gaze data to generate appropriate joystick values.

The floor/non-floor information is used to distinguish between driving and non-driving state. They coordinate of the gaze, which represents the height of the gaze point from the floor, is not used during the mapping, according to the fact that only fixations on the floor are translated into driving commands. This strategy is used to address the Midas Touch problem and is based on the results of the analysis of the participant natural gaze data.

## IV. Decoder development

### A. Continuous Control Field

The first interface uses a continuous control field, similar to those proposed by Ktena et al. [Sofia Ira Ktena 2015], Differently from the “Look Where You Want to Go” interface, the new proposed approach uses the actual distance of the user’s gaze point from the wheelchair (z coordinate), together with the X coordinate of the gaze, to generate the appropriate wheelchair command. Essentially, the two-dimensional control field is now projected on the ground and is represented by a rectangular portion of the floor, limited on the system FOV. Based on the results of the analysis of people’s gaze pattern, the floor area is divided into two different regions, for forward and backward motion. In the forward region, the equations, as functions of x and z, were used to predict the linear and angular velocities of the wheelchair, and were computed based on the analysis of the gaze data collected during the recordings. The objective was to find two fitting functions which predict, given the x and z coordinates of the gaze point on the floor, x and y joystick values as close as possible to the observed ones. Given the independent variables (x and z coordinate of the gaze point), the mean for every joystick values (dependent variable) associated with a specific gaze coordinate, has been used to find the two fitting functions. Polynomial functions of seven different degrees have been fitted to the data, and K-fold cross-validation (k=5) has been used to choose those providing the lowest Mean Square Error (MSE), on average (fig. 11). To do so, the data set is randomly divided into 5 different partitions. For each cycle of the 5-Fold cross-validation, the fitting functions are computed on 4 partitions. The MSEs are computed on the test dataset. In the 5 cycles, the one-fifth of samples, used as the test set, changes.

**Fig. 11.**
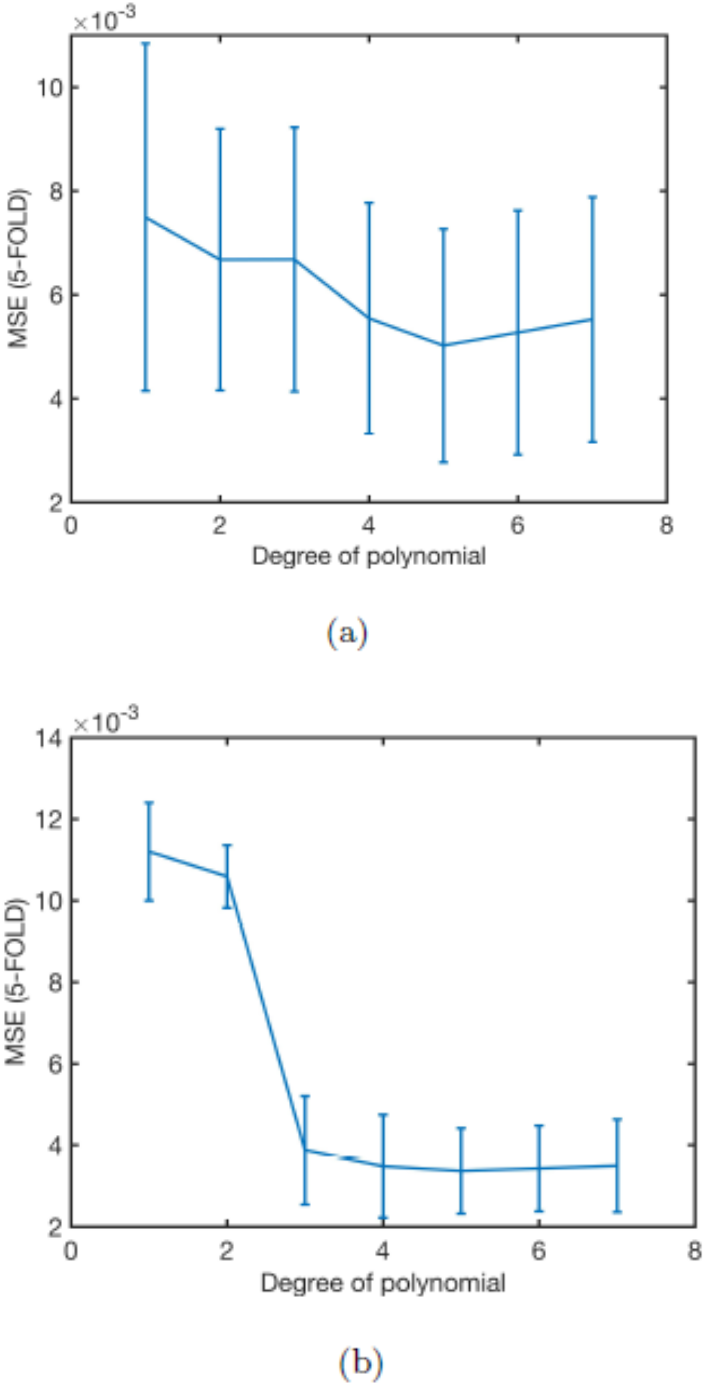
MSEs computed performing a 5-Fold cross validation for the fit of seven different polynomial orders. Error bar represent standard deviation over the 5 cycles. (a): MSEs for the fitting functions to map the x position of the gaze to angular velocity; (b): MSEs for the fitting functions to map the z position of the gaze to linear velocity.

The average MSEs over the cycles is then reported for each polynomial to choose the function with the lowest error and fit it to the whole set of available data samples.

For the linear velocity in the forward direction, we can observe that the magnitude is minimum at smaller depth from the wheelchair (z small) and smoothly increases, reaching its maximum at the top of the forward zone (Fig. 12 (a)). The further you look, the faster you move. This is consistent with the fact that when the target location that the user wants to reach is further away, the wheelchair has a longer distance to cover and therefore can achieve a higher speed.

**Fig. 12.**
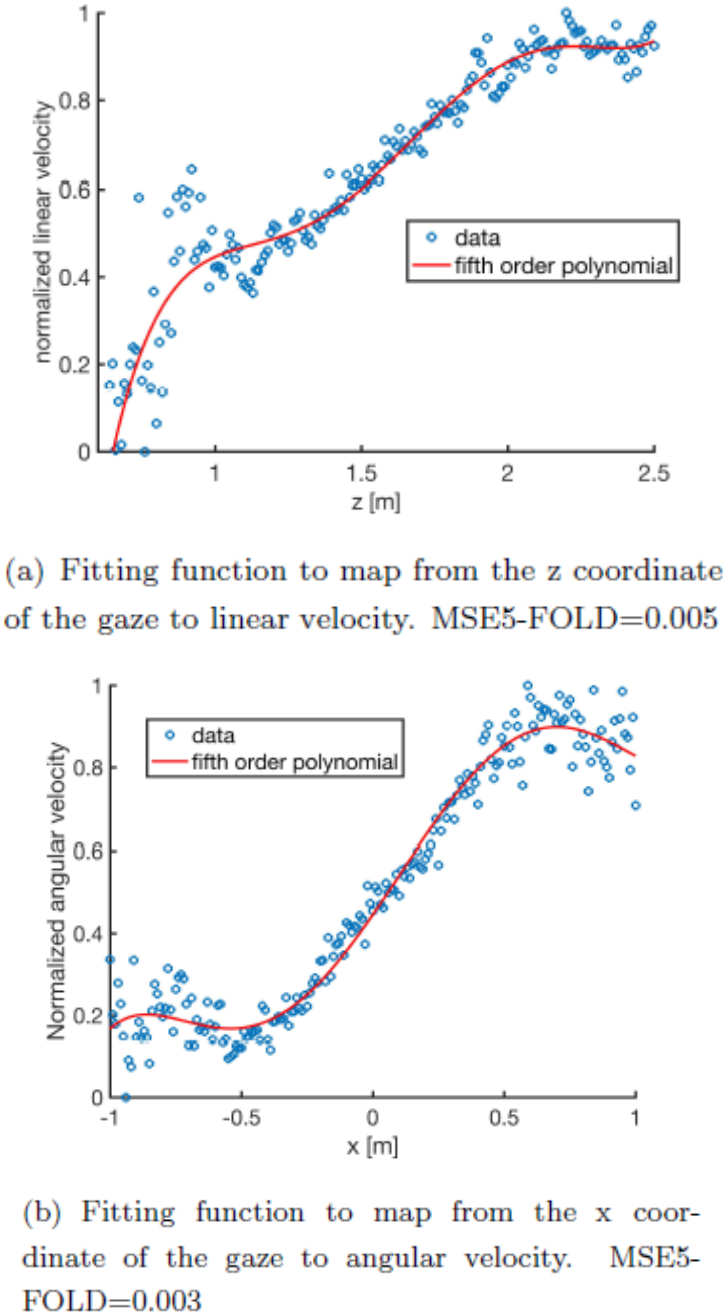
Fitting functions to calculate linear and angular velocity components for the wheelchair given the z and x coordinate of the gaze point on the floor during Forward Motion. The scatter plot represents the real data and the red line represent the best fit.

The fitting function for the angular velocity in the forward direction is shown in Fig. 12 (b). On the y-axis, the 0 value corresponds to the maximum angular velocity in the left direction, and 1 corresponds to the maximum in the right direction. We can observe that the angular velocity increases smoothly in the left direction when the user is looking further on the left. The same behavior can be seen in the right direction. Nevertheless, we can observe that at both extremities, the trend is different. This might be due to the instability of the joystick. In fact, the participants reported that they were unable to reach the extreme right directly and left joysticks directions. For this reason, the fitting function for the angular velocity is not reliable on the extremities. This will be taken into account during the controller implementation. The two fitting functions that derived from the user’s natural gaze behavior are scaled and shifted to produce values in the required range of the NI. The two equations are, therefore, used in the controller to generate a continuous control field similar to those shown in fig. 13. In the backward region, constant linear velocity is sent to the wheelchair, mainly for safety reasons.

**Fig. 13.**
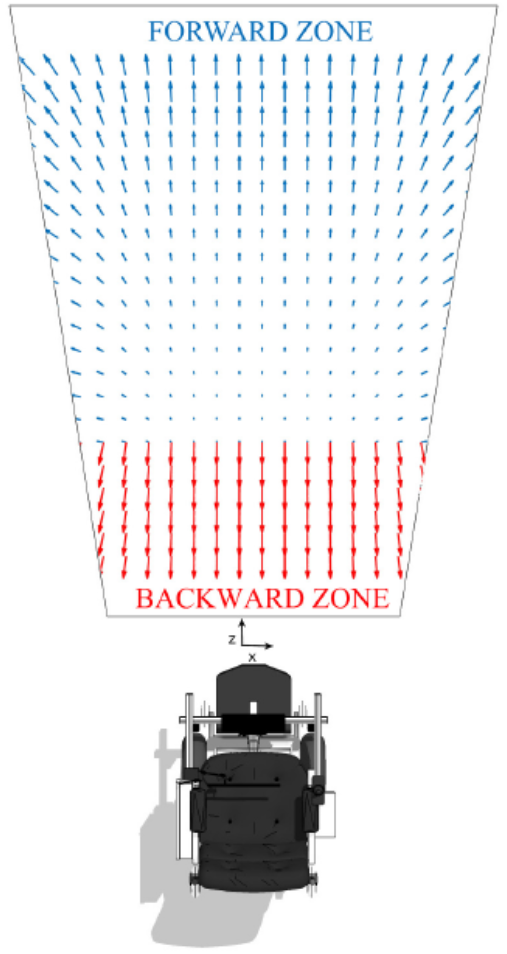
Continuous field control

### B. Natural Decoder

The second interface consists of a Natural Decoder, based on the analyzed people’s gaze distributions while driving. This decoder has the purpose of decoding real-time gaze data into wheelchair commands. To build such a decoder, eight different driving states *S_i_* (*i = 1, … 8*) have been identified: Straight Forward, Straight Backward, Forward Left, Forward Right, Backward Left, Backward right, Straight Left and Straight Right. Given the x and z coordinate of the gaze on the floor (*X_f_* = (*X_g_*, *Z_g_*)), the Maximum A Posteriori (MAP) decision rule is to apply to select the most likely driving state, according to the equation:

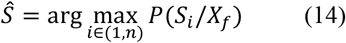

In (14), *P(S_i_/X_f_*) is the posterior probability that the user was in the state *S_i_*, given a gaze point of coordinate X_f_. The posterior probabilities are derived through Bayes’ rule:

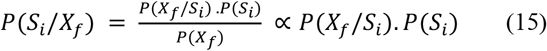

In (15), *P(X_f_/S_i_*) is the joint probability distribution of the user’s gaze point on the floor, during the state S_i_ and *P(S_i_*) is the probability of each state. For each state, the joint probability distribution is estimated based on the large data set collected during the subject recording. It is approximated as a 2D Gaussian, parameterized by a mean vector, n = (n1, n2), and a 2 x 2 covariance matrix C and with probability density:

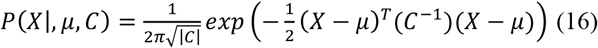

The parameters of the eight Gaussian distributions are computed by Maximum Likelihood Estimation (MLE). The probabilities *P(S_i_*) are computed based on the data collected during the experiments. They correspond to the probability that the user was in one of the eight driving conditions while navigating with the wheelchair. Finally, this set of parameters are implemented in the controller, based on the MAP decision rule of (14), to decode the real-time data of the driver into motor intention and generate the motion of the wheelchair accordingly. The posterior probability distributions involved in this rule, calculated using (15), are approximated as 2D Gaussian distributions, which are shown in fig. 14.

**Fig. 14.**
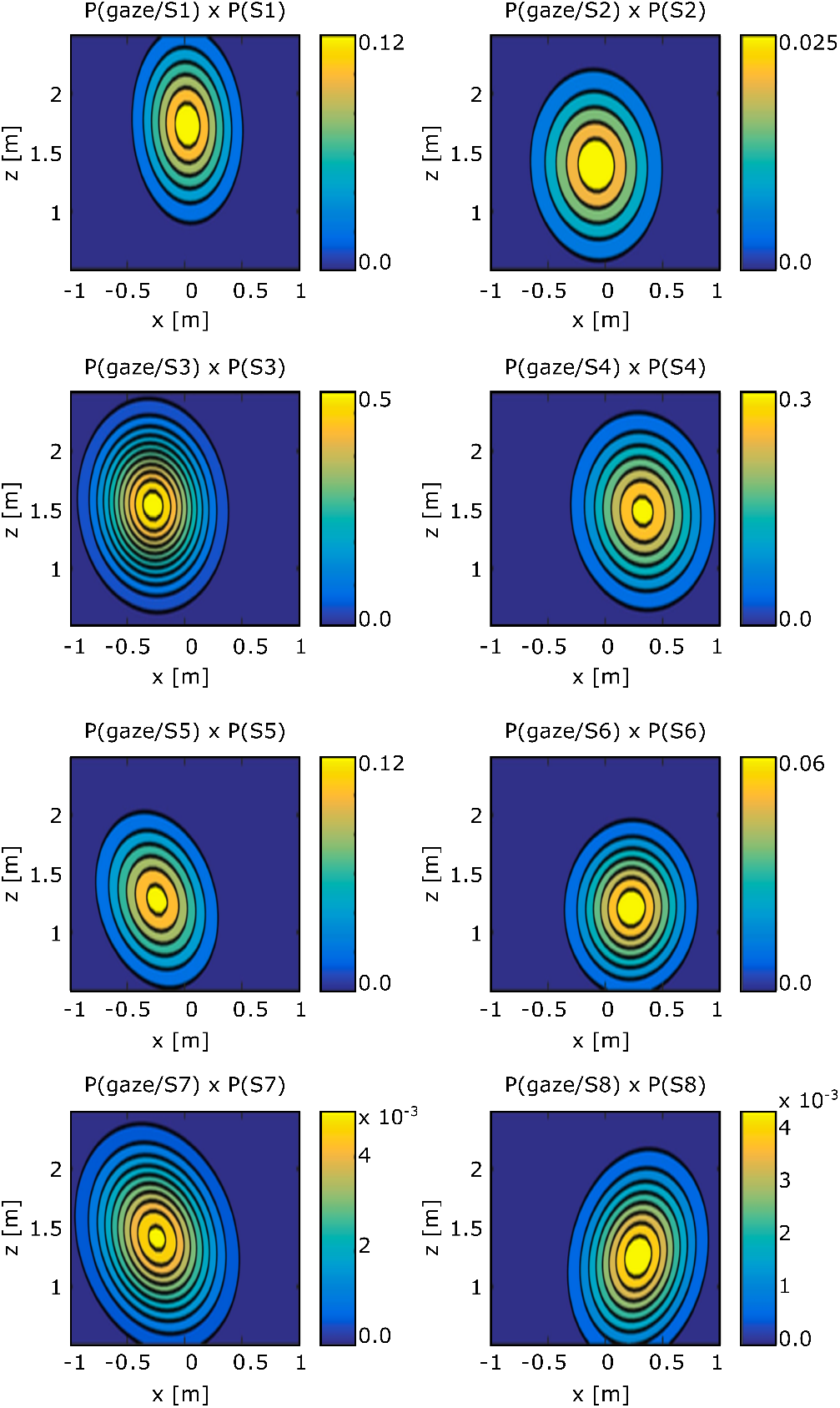
Posterior probability distributions of 8 different motion states approximately as Gaussian distributions. S1: Straight Forward; S2: Straight Backward; S3: Forward Left; S4: Forward Right; S5: Backward Left; S6: Backward Right.

To implement the natural decoder, for each gaze point on the floor, eight probabilities are computed using these distributions. Note that the range of the probabilities varies from one state to another. This is a question of visualization. In fact, the two last states present extremely low probabilities compare to the others. More precisely, for the case of the forward state (*P*(*S1|gaze*)) the center of the Gaussian distribution, is located inside the system workspace at high z values, and small x values. Therefore, when the user is looking in the forward zone, there is a high probability that the forward state will be selected. Similarly, this characteristic can be observed for the backward states, where the Gaussian is centered at smaller z values. Moreover, the center of the Gaussian for the motion states including right and left directions are slightly shifted to the right and the left, respectively. This was expected based on the results obtained in the analysis of the natural gaze pattern. To finally decode the user’s intention in real-time, the decoder selects the maximum probability over the eight distributions of fig. 14, given the user’s gaze point in the workspace. However, due to the very limited FOV of our system, the distributions are overlapping significantly. This means that only the states presenting a high range of posterior probabilities (Forward, Forward Right and Forward Left) can be selected when MAP decision rule is applied. Therefore, a small shift is introduced in the Gaussian distributions to reduce the overlap and allow the user to choose between a higher range of motion states. By associating each point in the workspace to the state which corresponds to the maximum probability over the 8 distributions, a colored map has been generated to visualize the effect of the natural decoder (fig. 15). Each shaded region corresponds to a specific motion state. We can observe that the straight right and straight left states are not present in the map. This is explained by the fact that the users rarely sent the extreme right and left commands, mainly due to inadequate joystick response for these directions. Hence, everywhere in the FOV, the posterior probability of these states is significantly lower than those of the other states and, therefore, are never selected. When focusing on one of these regions the user can steer the wheelchair in the corresponding direction.

**Fig. 15.**
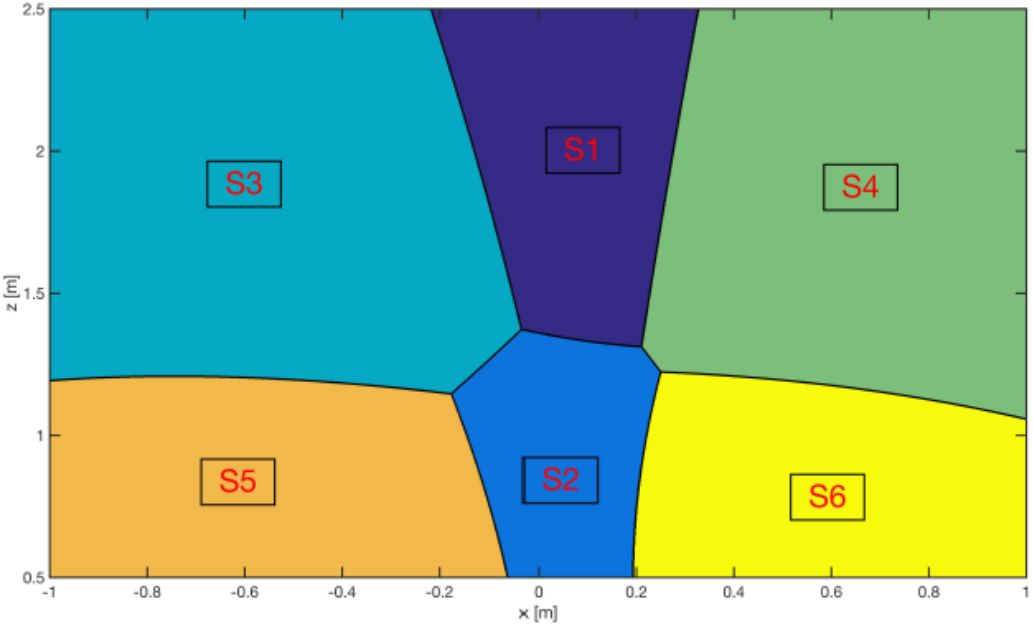
Colored map Natural Decoder. At each point in the workspace is associated the most probable state. Each color represents a particular driving state. S1: Straight Forward; S2: Straight Backward; S3: Forward Left; S4: Forward Right; S5: Backward Left; S6: Backward Right.

### C. Efficiency of the Wheelchair Controller Interfaces

Once the two above interfaces implemented, we tested their efficiency. Three subjects took part in this experiment and were naive to wheelchair use. Just one of them was wearing glasses. They were asked to complete three tasks by controlling the eye-based powered wheelchair with four different controller interfaces. We used four strategies for eye-based wheelchair navigation to compare the efficiency of our advanced user interfaces with strategies that have been previously implemented. The two controller interfaces for vision-based wheelchair navigation were taken to compare with our Natural Decoder Interface and Continuous Control Field Interface.

They are the following:

1. The Natural Decoder Interface.
2. The Continuous Control Field Interface.
3. The “Look Where You Want To Go” Interface. (**T**his interface was implemented using an NI Data Acquisition (DAQ) and developed by Sofia Ktena in [Sofia Ira Ktena 2015]. This interface is similar to the Continuous Control Field described above. Nevertheless, the gaze data are not expressed in metres, and the field of view is not on the floor. The x and y coordinates of the gaze point in pixels are translated into equivalent joystick inputs and send to the NI DAQ following an equivalent range of values.)
4. The Button Screen-Based User Interface. This interface has been implemented for the experiments. It is based on the screen-based interface presented in [Erik Wastlund 2010]. This interface corresponds to the vector field generated based on the button screen strategy and introduced in [Sofia Ira Ktena 2015]. The four limited regions corresponding to each button represent the motion states where the direction of the arrows indicates the wheelchair direction and the arrows length, the velocity of the wheelchair during motion. Outside these zones, there is no motion generated and is called the Idle state. For each zone the velocity is constant, and the values sent for respectively Left/Right and Forward/Backward commands are the following:(a) Forward: (128,190)(b) Forward Right: (190,190) (c) Forward Left: (80,190)(d) Backward: (128,80)

The three tasks (see Fig. 16) established during the experiments were:

1. Navigation along a free hallway
2. Room with obstacles: Slalom between 3 chairs and reach a table. Go back and slalom again between the 3 chairs.
3. Navigation along a free hallway to go back to the starting point

**Fig. 16.**
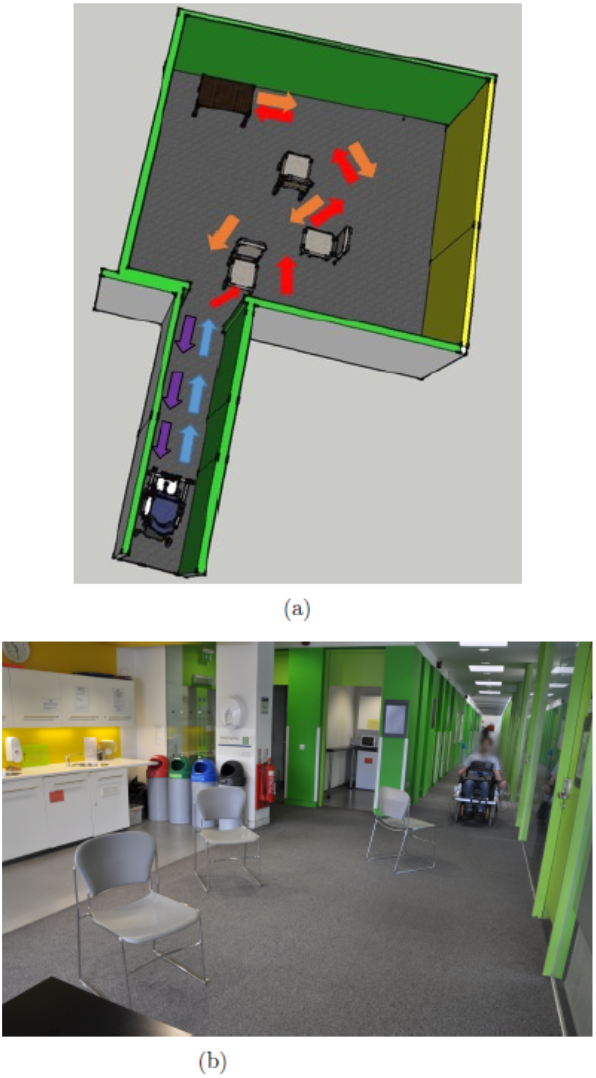
Experimental conditions a) Sketch of the circuit established for the experiments to test the efficiency of the controller interfaces. The blue arrows correspond to the circuit for the first task, the red and orange arrows for the second and the purple for the third task. b) Photo of a subject performing the tasks during an experiment.

The tasks were completed without any breaks to evaluate the efficiency of the controller for an entire exercise. Fig. 16(a, b) represents the sketch of the tasks and a picture taken during an experiment. For each task, different indicators were evaluated to allow the comparison of performance between the four interfaces.

1. The time of execution
2. The number of times that the user stops the wheelchair. This indicates the difficulty of finding the driving zones due to system limitation.
3. The number of times that the user uses the joystick instead of the gaze driven controller. This illustrates the difficulty of the certain task and the desire of using a more common interface instead of the gaze driven strategy.
4. The number of times the user pushes on the emergency button. This implies that the user encounters a situation requiring immediate stoppage of the wheelchair.

The efficiency of the above controller interfaces has been evaluated using specific indicators for each task completed during the experiments. Table 1 summarizes means of the results acquired for each controller interfaces and each task.

**Table 1.**
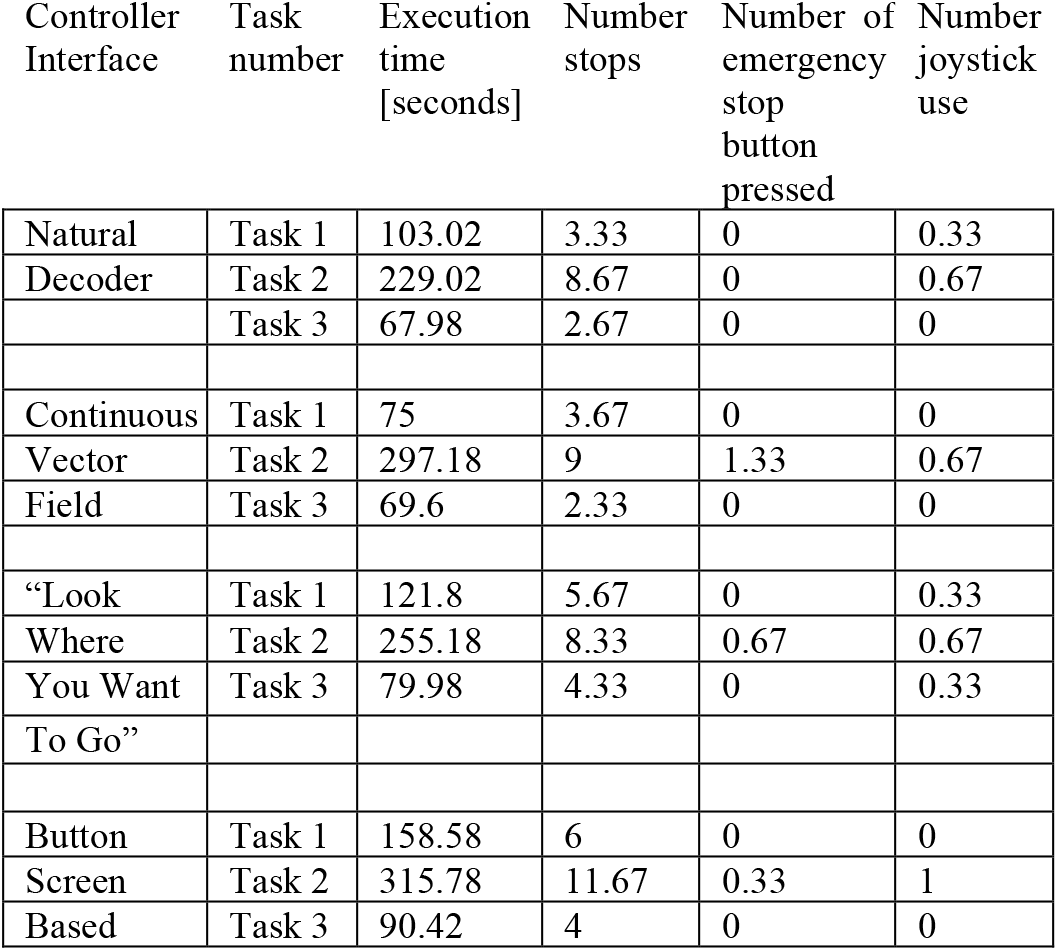
Results (mean) of the efficiency evaluation

| Controller Interface | Task number | Execution time [seconds] | Number stops | Number of emergency stop button pressed | Number joystick use |
| --- | --- | --- | --- | --- | --- |
| Natural Decoder | Task 1 | 103.02 | 3.33 | 0 | 0.33 |
|  | Task 2 | 229.02 | 8.67 | 0 | 0.67 |
|  | Task 3 | 67.98 | 2.67 | 0 | 0 |
| Continuous Vector Field | Task 1 | 75 | 3.67 | 0 | 0 |
|  | Task 2 | 297.18 | 9 | 1.33 | 0.67 |
|  | Task 3 | 69.6 | 2.33 | 0 | 0 |
| “Look Where You Want To Go” | Task 1 | 121.8 | 5.67 | 0 | 0.33 |
|  | Task 2 | 255.18 | 8.33 | 0.67 | 0.67 |
|  | Task 3 | 79.98 | 4.33 | 0 | 0.33 |
| Button Screen Based | Task 1 | 158.58 | 6 | 0 | 0 |
|  | Task 2 | 315.78 | 11.67 | 0.33 | 1 |
|  | Task 3 | 90.42 | 4 | 0 | 0 |

The fig. 17(a) represent the time of execution of the complete experiment averaged over all the subjects and fig. 17(b) is for the number of times the users stopped the wheelchair.

**Fig. 17.**
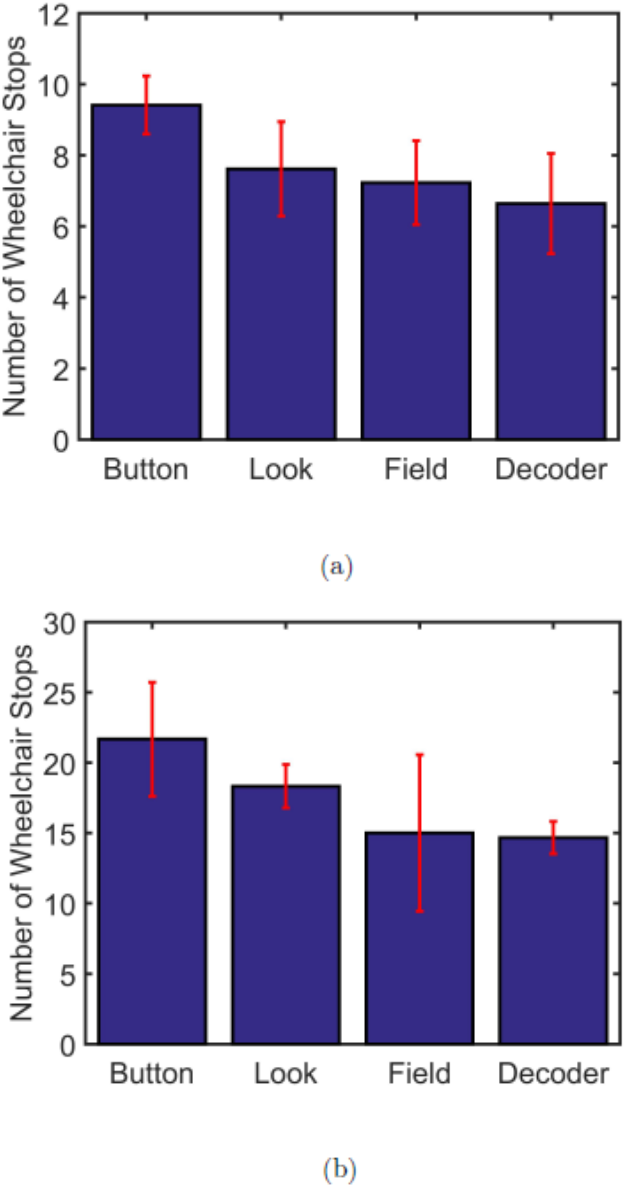
Efficiency Controller Interface. Button = Button Screen-Based [7]; Look = “Look Where You Want To Go” [28]; Field = Continuous Control Field; Decoder = Natural Decoder. Average and standard deviation over all the subjects for each interface a) of the time of execution over the entire experiment b) of the number of times the wheelchair stops over the entire experiment.

We observe that the execution time for the entire completed experiment is on average shortest whilst navigating the eye-based wheelchair with the natural decoder controller interface. This demonstrates that this strategy generates the most intuitive interface to control the powered wheelchair with an innovative mechanism. The continuous vector field developed for this project also shows better performance in term of execution time compared to the “Look Where You Want To Go” and the Button Screen Based interface. More precisely, subjects seemed to show more facility during the second task with the natural decoder. This task had a higher level of difficulty due to the presence of several obstacles. This proves that the controller interface shows more potential when the gaze driven controller is targeted towards the floor level, and when this interface is decoding natural gaze behavior.

It can be observed in Fig. 17 (b) that the number of times that subjects stopped the wheelchair was on average lower for the two developed interfaces, compared to the “Look Where You Want To Go” and the screen-based button interface. This means that even with a limited working space due to the entire system restricted field of view, the subjects still managed to find the driving zone easily. This driving zone is characterized by a working space on the floor level which indicates that this property is significantly present in natural gaze behavior during vision-based wheelchair navigation. Furthermore, during the experiments, the user was able to look around and talk to people surrounding him without generating wheelchair motion. This is due to this natural behavior directed towards the defined driving zone, which mainly illustrates drastic reduction of the Midas Touch problem.

Furthermore, it can be observed in Table 1 that while performing the tasks with the natural decoder, none of the subjects needed to use the emergency button to produce immediate stoppage of the wheelchair. They had a significant amount of control over the eye-based wheelchair while using the natural decoder interface. As concluded in [Al Haddad et al., 2011], providing the user with more control over the driving wheelchair leads to an efficient system and increases user satisfaction.

Finally, due to the low number of stops and the short execution time, the natural decoder shows excellent performance even when the user wanted to go backward. They were able to distinguish the forward from the backward in a natural way. The natural decoder has improved the controller efficiency for this particular case and increased the user interface intuitiveness.

Nevertheless, Table 1 also shows that subjects seemed to perform faster the task in the free hallway with the continuous vector field. This can be explained by observing the fig. 15 In the set of state conditional probabilities in the natural decoder, the motion state probabilities corresponding to the straightforward command are delimited by a small region reducing the chance of going straight forward and increasing the probability of deviating to the right and left side. In conclusion, in a free hallway, where the main motion state is straightforward, controlling the eye-based wheelchair with the continuous vector field is easier than with the natural decoder. In summary, the natural decoder developed for this project provides a user-friendly and intuitive interface for the eye-based powered wheelchair navigation using natural eye movements. This interface increases the user’s confidence by giving substantial control over the system. Finally, it allows the user to drive in a dynamical environment. The natural behavior has been shown high efficiency during the experiments. The floor/non-floor attributes to the motion/no motion states reduce the Midas Touch problem drastically and allow the user to look around without leading the wheelchair into motion.

## V. Discussion

A 3D gaze-tracking system has been designed, with the primary goal of analyzing people natural eye movements in the context of wheelchair navigation. The system uses two low-cost devices: a remote eye-tracker, and an RGB-D camera, to estimate the gaze point of the user in a 3D environment.

The depth sensing technology of the RGB-D camera is used to translate the two-dimensional information provided by the eye-tracker into three-dimensional gaze data, and to provide information to distinguish between gaze points on the floor or not on the floor. Experiments have been conducted to test the accuracy of the 3D gaze estimation. The system performs with a mean Euclidean error of 16.7 ± 3.3 cm (mean ± standard deviation). This corresponds with the level of performance in comparison with other low-cost devices (less than $100 each) [Abbott and Faisal 2012], Moreover, in the context of analysis of gaze patterns during wheelchair navigation, this level of accuracy can and should be considered completely acceptable.

The main finding resulting from analysis of people’s gaze patterns was that the wheelchair’s drivers tended to direct their fixations toward the floor more often when they navigated in regions cluttered with obstacles, than when they steered in a free environment. Intuitively, when the complexity of the situation increases, the individuals need to look at the floor more often. This might be explained by the fact that, in a clear of obstacles environment, people can detect the absence of obstructions with peripheral vision and, therefore, do not require to consistently locate the gaze downward [Mark A Hollands 2002]. On the other hand, while navigating in the rooms with obstacles, people need to look at the floor more often, to identify the path to follow and reach the targeted location, avoiding collisions with the objects.

We found that the drivers had a preference to direct their fixations on the floor while moving with the wheelchair. This finding is consistent with previous results by Einhäuser W, 2007. Authors showed that subjects made more likely down-vertical oriented eye-movements when they were walking in an obstacle-filled or less predictable environments, e.g. at the furnished apartment or in a forest road full of puddles [Einhäuser W, 2007]. In our study, subjects tended to focus their visual attention on the path they wanted to follow, in accordance with what has been found in previous work [Paul L Kaufman et al. 2011]. This finding had an essential impact on the strategy for the eye-based controller interfaces proposed in this project. It helped us to make a distinction between driving and the idle state, drastically reducing the effect of the Midas Touch problem, naturally and intuitively.

The second main finding of this analysis was that the users tended to look further away on the floor from the wheelchair while moving forward and closer to the wheelchair while moving backward. Furthermore, in the forward motion, there was a clear preference for the participants to look right or left, respectively, while moving in the right and left direction, respectively. Therefore, it is possible to state that the participants, while driving a wheelchair, tended to look in the direction where they are moving. This is consistent with previous studies, which showed the existence of such a gaze pattern in other various forms of locomotion, from walking [Takahiro Higuchi 2013] to driving a car [Takahiro Higuchi 2014]. However, a general pattern could not be identified for the backward right and backward, left motion. This was because the participants showed different behaviors: some users tend to look on the right while moving in the backward right direction, others showed the opposite gaze pattern.

Based on the analysis of people’s gaze patterns, two different interfaces were developed. The “Continuous control field interface,” is based on natural eye movements, in that the equations used to predict the linear and angular velocity of the wheelchair have been estimated from the data collected during the recordings. The “Natural Decoder,” on the other hand, is directly built on the drivers’ gaze distributions while moving in specific directions. These distributions are used to decode realtime gaze data into the motor intention of the driver.

While testing on different subjects, both the interfaces proved to be intuitive and user-friendly. The “Natural Decoder” showed better performance, especially while navigating in more complex environments. Using this interface, it was easy to move around obstacles, with a good control of the chair. Control of backwards motion, which represented one of the main drawbacks of “Look Where You Want To Go” interface proposed by Ktena et al. [Sofia Ira Ketna 2015], was intuitive and easy to select using the “Natural Decoder.” On the other hand, the “Continuous Control Field Interface” was preferred when the users had to follow a straight trajectory. Using the “Natural Decoder,” the drivers were often not able to select the straight ahead motion and often deviated to the right and left directions. This is explained by the fact that the floor region, where the user has to look to select the forward command, is quite limited in size, as can be seen in fig.15. During free navigation, the participants tended to move right forward and left forward and rarely drive in a straight direction. Both the interfaces generate the motion of the wheelchair only when the gaze point of the user is on the floor. This strategy, which is based on the natural tendency of people to look at the floor while moving, was used to make a distinction between motion relevant and motion irrelevant eye movements. Through this, we proposed a solution, for a gaze-driven wheelchair, which can limit the effect of the Midas Touch problem, avoiding other gestures, such as blinking and winking, from the users.

As previously mentioned, the designed 3D eye-tracking system reaches a satisfying level of performance, especially when considering the low cost of the whole hardware setup. However, it does show several limitations, with disadvantages for both the analysis of natural gaze behavior and the implementation of the controller interface. First of all, one of the primary constraints of our study has been the size of the FOV of the system. This is due to the limited FOVs of both eye tracker and depth camera. As a consequence of this, the gaze of the participants could be tracked only in a limited space area. Furthermore, the reliability of the data, used to investigate people’s gaze patterns, is further limited by other two factors. Firstly, the 3D gaze estimation error is considerably dependent on the depth. This makes gaze data, also estimated from the wheelchair, less reliable. Secondly, with the proposed system, the eye tracker calibration cannot be verified quantitatively. Therefore, during the recordings, a potential loss of the calibration could only be checked qualitatively, reducing the reliability of the data and increasing the experiment duration. Ideally, the system should be able to automatically verify the accuracy of the calibration, to make sure that the gaze estimation error is always under an acceptable limit. This is particularly relevant in our application, where the system is mounted on a wheelchair and used for long lasting experiments, which results in a frequent loss of calibration.

